# Induction of meningioma stem signature *via* hydrogel reprogramming and application of meningioma stem cell marker CXCR4 to pathological diagnosis and treatment

**DOI:** 10.1101/2025.06.14.659672

**Authors:** Yoshitaka Oda, Masumi Tsuda, Lei Wang, Jun Suzuka, Sayaka Yuzawa, Kouki Ise, Umma habiba, Jintao He, Satoshi Tanikawa, Hirokazu Sugino, Zen-ichi Tanei, Christian Mawrin, Jian Ping Gong, Shinya Tanaka

**Affiliations:** Department of Cancer Pathology, Faculty of Medicine, Hokkaido University, Sapporo, Japan; Institute for Chemical Reaction Design and Discovery (WPI-ICReDD), Hokkaido University, Sapporo, Japan; Department of Diagnostic Pathology, Asahikawa Medical University Hospital, Asahikawa, Japan; Department of Surgical Pathology, Hokkaido University Hospital, Sapporo, Japan; Department of Neuropathology, Otto von Guericke University Magdeburg; Faculty of Advanced Life Science, Hokkaido University, Sapporo, Japan

**Keywords:** cancer stem cell, reprogramming, meningioma, double-network hydrogel, PNaSS gel, PCDME gel, soft matter, CXCR4

## Abstract

**Backgrounds:** Meningioma accounts for about 40% of brain tumors, but there was no effective treatment for recurrent or inoperable cases. We have previously found that when cancer cells are cultured on certain hydrogels, cancer stem cells are efficiently induced in various cancer types and we named this process as hydrogel activated reprogramming (HARP) phenomenon. In this study, we aimed to identify a key molecule that induced meningioma stem cells *via* hydrogels.

**Methods:** Meningioma cells cultured on hydrogels were analyzed for expression of regular stem cell markers and their tumor genericity. Microarray analysis was performed to identify meningioma stem cell specific markers and to examine the application of such marker molecules as a therapeutic targets or pathological diagnosis for grading.

**Results:** Regular stem cell markers such as *Nanog*, and *Oct3/4* were induced by culturing meningioma cells on hydrogels, and comprehensive gene expression analysis identified molecules involved in cancer stem cell activity. Among them, CXCR4 was selected as a therapeutic target molecule. Stimulation of CXCR4 *via* CXCL12 led to an increase in stem cell markers. In human meningioma pathological specimens and cultured cell lines, there was a correlation between CXCR4 expression levels and NF2 mutations and/or deletions. CXCR4 immunohistochemistry also detected along with the brain invasion. Thus, CXCR4 immunohistochemistry may be useful to suggest typical CNS WHO grade 1 meningioma, that do not require molecular analysis.

**Conclusions:** We have defined meningioma stem cell signature *via* HARP phenomenon and identified CXCR4 with biological significance as being diagnostic target.

**Key Points:**

- We induce meningioma stem signature *via* Hydrogel activated reprogramming (HARP) phenomena.
- We identified CXCR4 as a candidate molecule as stemness marker and therapeutic target.
- Translational research using pathological specimens reveal CXCR4 immunohistochemistry has been shown to play an auxiliary role in the diagnosis of meningiomas.

**Importance of the Study:** In addition to morphology, immunohistochemistry and gene alteration are being adopted as diagnostic criteria for CNS tumors. From CNS5 onwards, epigenetic changes such as methylation analysis are being adopted as diagnostic criteria. We induced epigenetic changes in meningioma cells and induced cancer stemness, and this technique will be of great importance in the research and diagnosis of meningioma. In addition, microarray analysis was used to select CXCR4 from the molecules that increased simultaneously during stem cell induction on the three hydrogels, and further analysis revealed that CXCR4 immunostaining may indicate the distribution of meningioma stem cells, also making it useful for diagnosis. This report will contribute to the advancement of meningioma research and diagnosis by conducting basic research and using histopathological specimens.

## Introduction

Meningioma accounts for about 40% of brain tumors ^1^. Although many patients are operable and have a better prognosis than other brain tumors, meningiomas classified as WHO grade 1 have recurrences within about 10 years of about 15-20%^2^. The recurrence rate of WHO grade 2 and WHO grade 3 is high, at least 50%^34^. Although radiation therapy shows some effect for inoperable cases, no effective drug treatment has been established^5^. Gene mutations that are associated with poor prognosis such as deletion and mutation of NF-2 and mutation of the TERT promoter region, and those that have good prognosis such as TRAF7, AKT1, KLF4, and SMO have been identified ^67^. However, the mechanisms of tumorigenesis and treatment resistance have not been elucidated, and no effective therapeutic target molecule has been identified.

On the other hand, it has been revealed that tumor cells are not uniform and that cancer stem cells present in a part of the tumor are greatly involved in tumor recurrence and treatment resistance ^8^. Eradication of tumors requires treatment for cancer stem cells, but the number of cancer stem cells in tumor tissue is small, and a method for efficiently inducing cancer stem cells has not been established. Nanog, Oct3/4, and Sox2 are representative markers of pluripotent stem cells. In meningiomas, Nanog, KLF4, Oct3 / 4, CD44, CD133, Sox2, Nestin, Frizzled 9 and others have been reported to be involved in cancer stem cell ^9,10,11^, except for Nanog, there are variations depending on the report.

We have previously found that when cancer cells are cultured on certain hydrogels, cancer stem cells are efficiently induced in various cancer types ^12–15^. Double-network hydrogels (DN gels) invented by Gong *et al*. are soft and tough materials containing 85∼90wt% water ^16,17^. Due to its similarity to biological tissues that are also soft and water-containing materials, DN hydrogels show unique bio-properties, such as inducing the regeneration of cartilage in the knee joint of rabbits^18^. The PAMPS/PDMAAm DN gels have a big potential to produce CSCs from non-CSCs rapidly and efficiently within 24 hours, which is valuable to analyze the CSC’s characteristics and identify anti-cancer drug that selectively eradiate CSCs19.

The PCDME gel (poly (N-(carboxymethyl) -N, N-dimethyl-2-(methacryloyloxy)) gel) led to the induction of cancer stemness in pancreatic cancer cells. ^14,20^. The PNaSS gel is poly (sodium p-styrene sulfonate) and we demonstrated that PNaSS gel is involved in the induction of cancer stem cells in leukemia cells, and identified candidate therapeutic molecules through single-cell analysis^15^.

In this study, we aim to identify key genes that induce meningioma stem cells via hydrogels, which might be an effective candidate therapeutic molecule for meningioma.

## Results

### Stemness nature was induced by culturing meningioma cells on hydrogel

One meningioma cell line (HKB-MM) and three primary culture cells (WY02, WY08, and WY09) were cultured on DN gel, PCDME gel, and PNaSS gel using conventional culture medium (Fig. 1*A*). On the DN gel and the PCDME gels, all meningioma cells formed spherical agglomerates, whereas the adhesive morphology was observed on the PNaSS gel as on the PS dish (Fig. 1*B*). The expressions of stemness markers *Nanog* and *Oct3/4* were up-regulated in all meningioma cells on three kind of hydrogels against PS dish, especially high on the PNaSS gel (Fig. 1*C*). In the WY02 cells, the expression of *Nanog* increased on all three hydrogels, but *Oct3/4* was elevated only on PNaSS gel (Fig. 1*C*). *SPP1* is a gene with the highest expression change in the DN-gel induced CSCs, of which expression was enhanced on the DN gel and the PCDME gel, but not on the PNaSS gel (Fig. 1*C*).

**Fig. 1.**
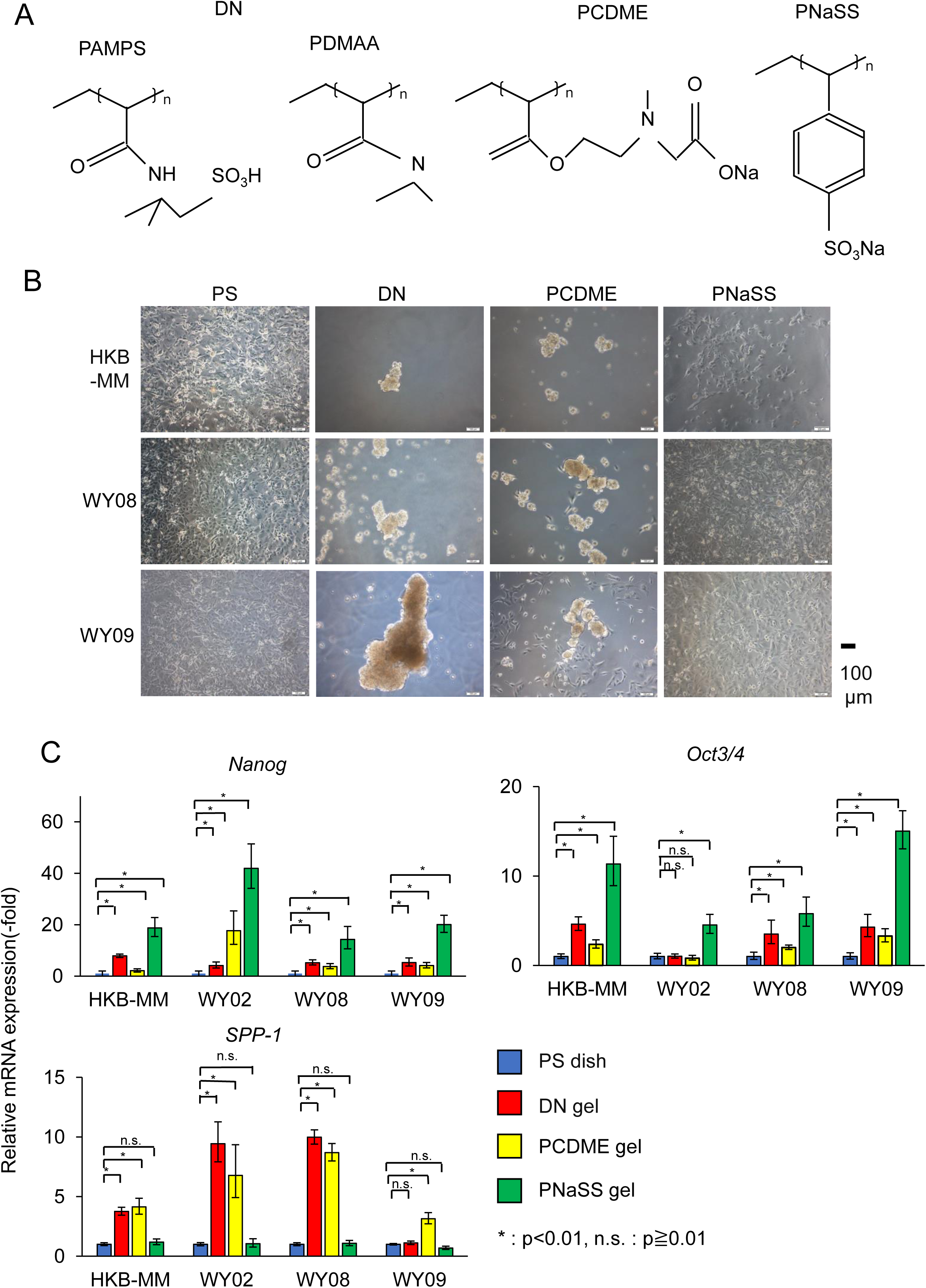
Morphological change and stemness induction on hydrogels in meningioma. **A**, Chemical formulae of DN gel components, poly(2-acrylamido-2-methyl-1-propane-sulfonic acid) (PAMPS) and poly(N,N-dimethylacrylamide) (PDMAAm), PCDME gel and PNaSS gel. **B**, DN gel and PCDME gel induced cell aggregation and forming spherical structures in one cell line (HKB-MM) and three primary culture cells (WY02, WY08, WY09). The adhesive morphology was observed on the PNaSS gel as on the PS dish. Scale bar = 100 μm. **C**, Real-time PCR analysis of mRNA levels of stemness markers (*NANOG*, *OCT3/4*) and *SPP-1* in four cells cultured on PS dish, DN gel, PCDME gel and PNaSS gel. Data are represented using two-sided unequal variance *t*-tests in comparison with the expression of PS. * :*P* < 0.01 vs. PS.

### CXCR4 was selected as a candidate molecule for treatment by comprehensive analysis

Here, we focused on WY08 and WY09 cells because of universal reactivity to all three hydrogels in the expression of *Nanog* and *Oct3/4* (Fig. 1*C*). Sequential cell culture revealed that the expression of *Nanog* and *Oct3/4* was most enhanced on the DN gel and the PCDME gel for 3 days, whereas on the PNaSS gel for 7 days Fig. 2*B*). To identify novel molecules associated with stem cell nature of meningioma, gene expression profiles were investigated by microarray analysis using samples extracted from WY08 and WY09 cells cultured on the DN gel and the PCDME gel for 3 days, and the PNaSS gel for 7 days (Fig. 2*A*). Under these situation, WY08 cells appear to be most responsive to the DN gels and WY09 cells to the PNaSS gel (Fig. 2*B*). In each WY08 and WY09 cells, we picked up genes that were more than 4.0-fold up-regulated on the DN gels, the PCDME gels, and the PNaSS gels compared to the PS dish, by using standardized data and represented them in a Venn diagram (Fig. 2*C*). There were 177 and 99 genes in WY08 and WY09 cells, respectively, which were commonly up-regulated on the three types of gels (Fig. 2*C*). Among them, 20 genes were commonly more than 4.0-fold up-regulated between WY08 and WY09 cells (Fig. 2*D*), and we selected CXCR4 as a candidate gene of therapy target by pathway analysis using DAVID. To confirm the elevation of *CXCR4* and its ligand *CXCL12*, we checked mRNA level of them by Real-time PCR. We confirmed that expression of *CXCR4 V1*, *CXCR4 V2*, and *CXCL12* were up-regulated in three meningioma cells on the DN gel, the PCDME gel, and the PNaSS gel (Fig. 2*E*). *CXCR4* and stemness markers such as *Nanog* and *Oct3/4* were enhanced with the treatment of SDF-1(CXCL12) recombinant protein in WY08 cell (Fig. 2*F*).

**Fig. 2.**
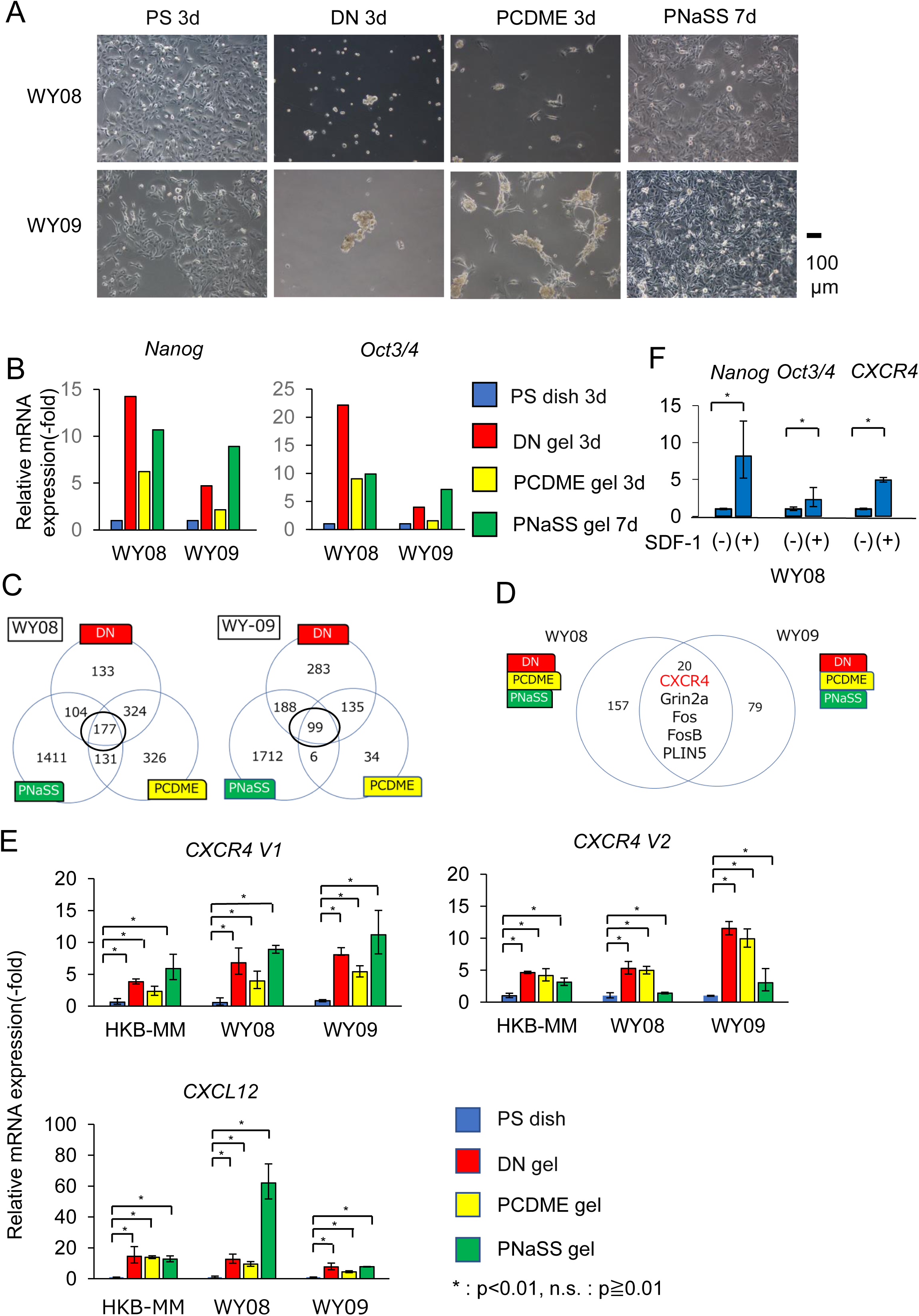
Microarray analysis: CXCR4 was increased in meningioma cells on hydrogels. **A**, Representative photographs of WY08 and WY09 cells cultured on PS dish in 3 days, DN gel in 3days, PCDME gel in 3days and PNaSS gel in 3days. Scale bar = 100 μm. **B**, Real-time PCR analysis of the mRNA levels of stemness markers such as *NANOG* and *OCT3/4* of WY08 and WY09 cells. **C,D**, Microarray analysis of WY08 and WY09 cells cultured on DN gel, PCDME gel and PNaSS gel. Venn diagram indicates the number of genes that up-regulated more than 4 times compared to those cells in normal culture condition on PS dish. **E**, Real-time PCR analysis of the mRNA levels of *CXCR4* in three cells (HKB-MM, WY08 and WY09) on DN gel, PCDME gel and PNaSS gel. The data are presented as the mean ± SD of three same experiments in triplicate, and represented using two-sided unequal variance *t*-tests in comparison with the expression of PS. * *P* < 0.01 vs. PS. f, Real-time PCR analysis of the mRNA levels of CXCR4 and stemness markers such as *NANOG* and *OCT3/4* of WY08, treating with SDF-1(CXCL12) recombinant protein 100 ng/ml.

### PNaSS gel increased *in vivo* tumorigenicity of meningioma cells

To evaluate the effect on tumor development *in vivo* in hydrogel-induced stem cells, we conducted experiments using a xenograft model of subcutaneous transplantation of WY08 cells with different cell number of 1.0 × 10^3^, 1.0 × 10^4^, and 1.0×10^5^ (n=4, 3, and 3). The cells cultured on the PNaSS gel acquired higher tumorigenicity than cells cultured on PS dish (Fig. 3*A*). Satellite nodules of the tumor were observed in 7 of 10 mice injected the cells cultured on the PNaSS gel, whereas which were observed only 1 of 10 mice in the case of PS dish (Fig. 3*B*, 3*C*). Venous invasion was observed in 3 of 10 mice injected the cells cultured on the PNaSS gel, whereas no invasion was observed in mice injected the cells cultured on the PS dish (Fig. 3*C*, 3*D*). In immunohistochemistry, CXCR4-positive cells were observed not only in tumor center but also at the invasive front in the tumors of mice injected the cells cultured on the PNaSS gel. Meanwhile, the staining at the invasive front was undetected in mice injected cells cultured on the PS dish (Fig. 6*E*).

**Fig. 3.**
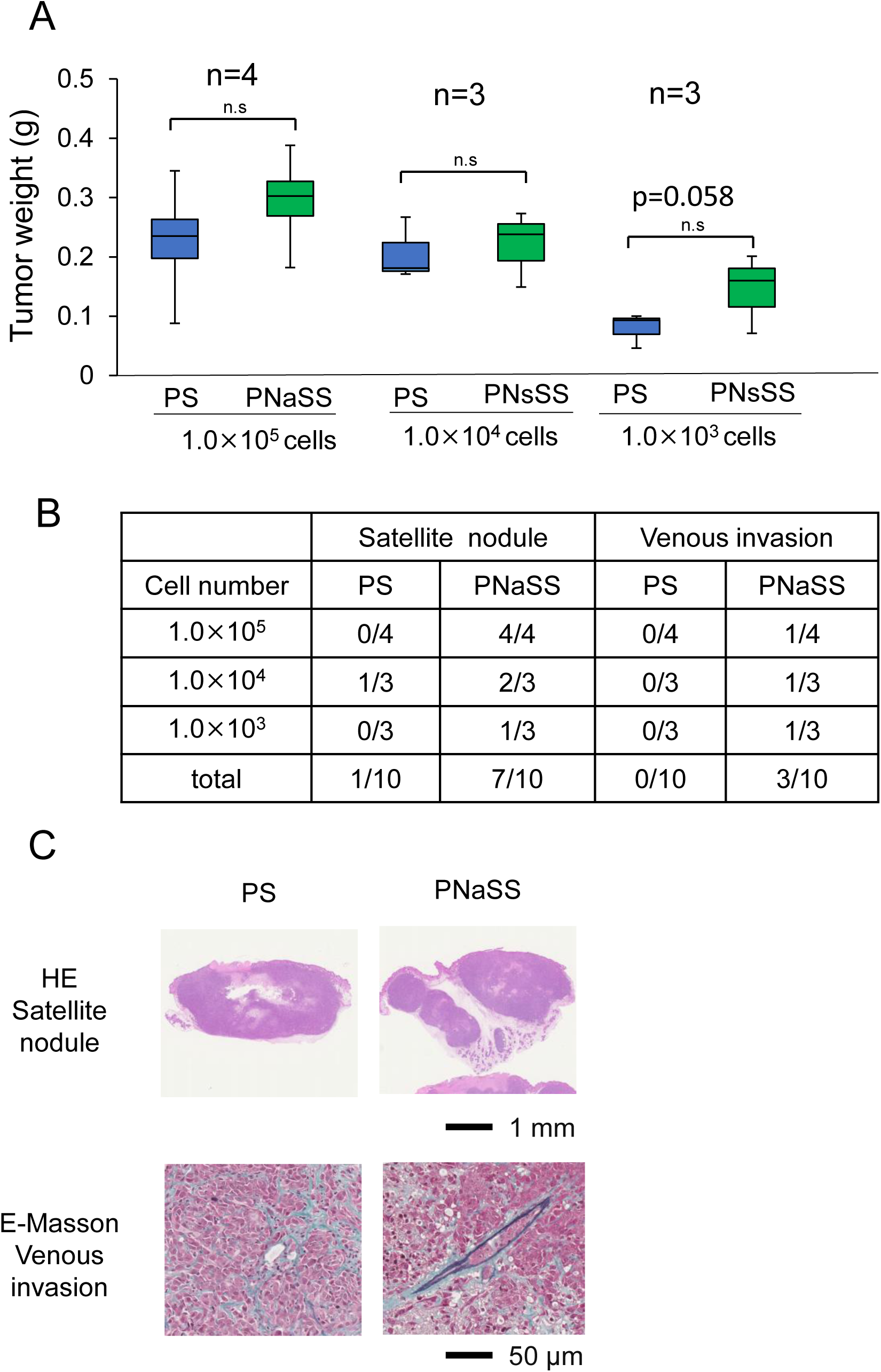
Xenograft model: PNaSS gel increased *in vivo* tumorigenicity of meningioma cells. **A**, Cells (WY08) were cultured on PNaSS gel or PS dish for one week and inject into subcutaneously in both abdomen. Left side(control): Cells cultured on PS dish. Right side: Cells cultured on PNaSS gel. The mice were sacrificed at 11 weeks. The number of cells were 1.0 x 10^5^, 1.0 x 10^4^, 1.0 x 10^3^. **B,** The tumorigenicity tended to be higher on the side injecting the cells cultured on the PNaSS gel than on the PS dish. **C,** Cells cultured on PNaSS gel formed more satellite nodules and more venous invasion than cells cultured on PS dish. **C,** Representative picture of satellite nodule (Right, PNaSS gel) and simple nodular tumor (Left, PS dish).

### Pathological specimen of Human meningioma

To investigate the pathological implications of CXCR4 in meningioma, CXCR4 expression in meningioma tissues was examined. The backgrounds of 40 meningioma patients used in the analysis are shown in Table 1. Staining of the vascular endothelium was settled as endogenous control. In the tumor tissues, the staining of CXCR4 was observed diffusely, as cluster with more than five cells, and in single cell, which were scored 3+, 2+, and 1+, respectively (Fig. 4*A*). Here, we investigated a correlation between immunohistochemistry for CXCR4 and NF-2 status, and for CXCR4 status and histological status. The 40 meningioma patients were classified histologically meningothelial type (17 cases) and other (23 cases) (Fig. 4*B,* 4*C,* 4*E*). Among them, the patients with NF2 mutation/deletion were 24 cases, which were dominant in patients with histological types other than meningothelial type (21/24), and the staining of were 14 and 10 cases for positive and negative, respectively (Fig. 4*B,* 4*D,* 4*F*). Conversely, the majority of patients without NF2 mutation/deletion (14/16, 87.5%) were of the meningothelial type (Fig. 4*B,* 4*D,* 4*F*). Thus, the patients with CXCR4 staining tended to be with NF2 mutations/deletion and histologically except for meningothelial type.

**Fig. 4.**
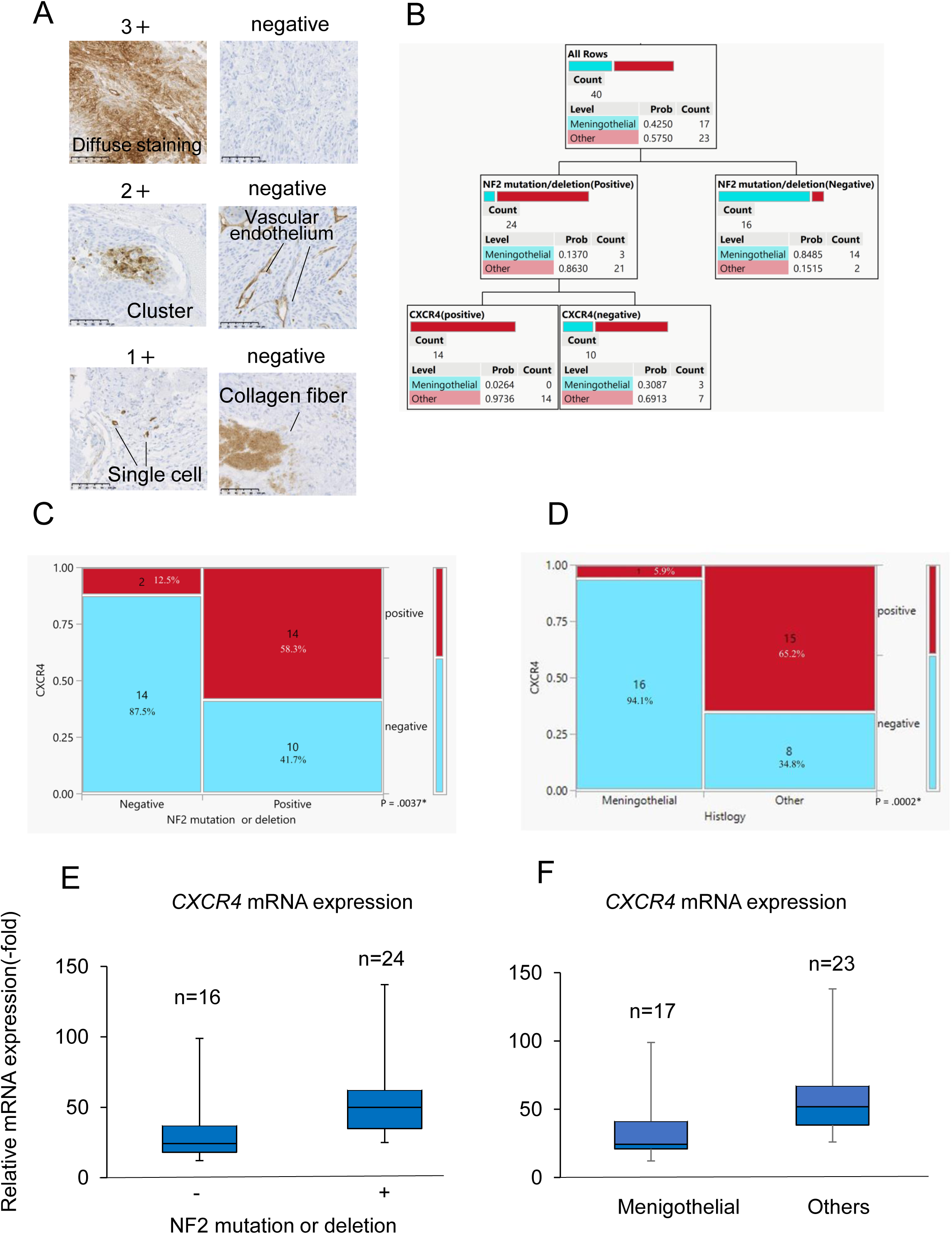
CXCR1 expression correlates with the malignant potential of meningioma. A-D, Immunostaining of CXCR4 in tumor specimens of 40 human meningioma patients are displayed. **A**, 0: Negative in all cells, Staining to the vascular endothelium or collagen fiber. 1: Weak staining compared to vascular endothelium. 2: Collection of 5 or more cells showing intensive staining compared to vascular endothelium (cluster formation), 3: Diffuse intensive staining compared to vascular endothelium. H&E staining was also performed. **B,** Flowchart for meningioma diagnosis including CXCR4 immunohistochemistry **C**, Comparison of CXCR4 immunohistochemistry with NF-2 mutation or deletion in human sample. **D**, Comparison of CXCR4 immunohistochemistry with histology in human sample. **E**, Comparison of CXCR4 expression with NF-2 mutation or deletion in human sample **F**, Comparison of CXCR4 expression with histology in human sample.

**Table 1.**
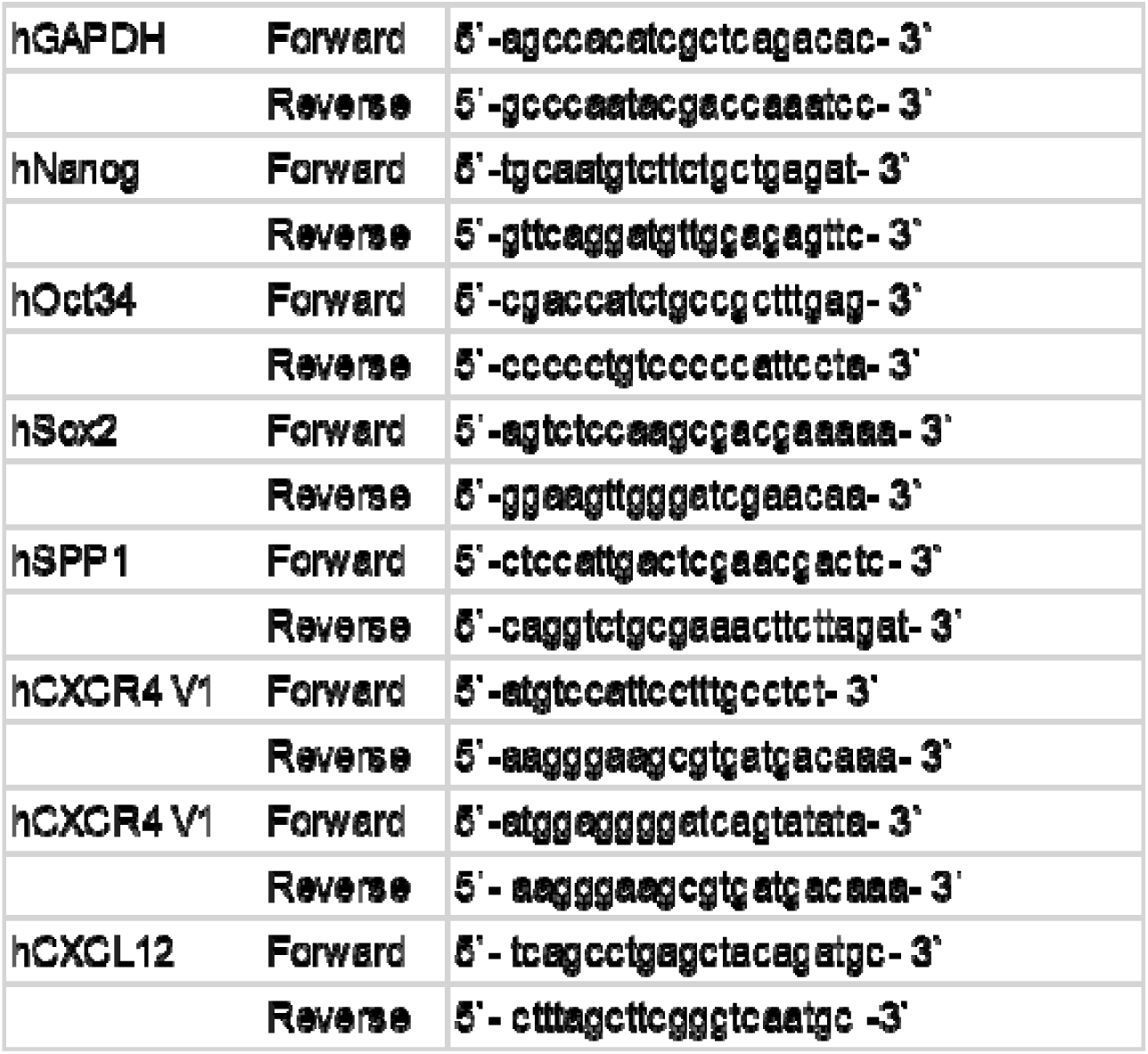

**Table 2.**
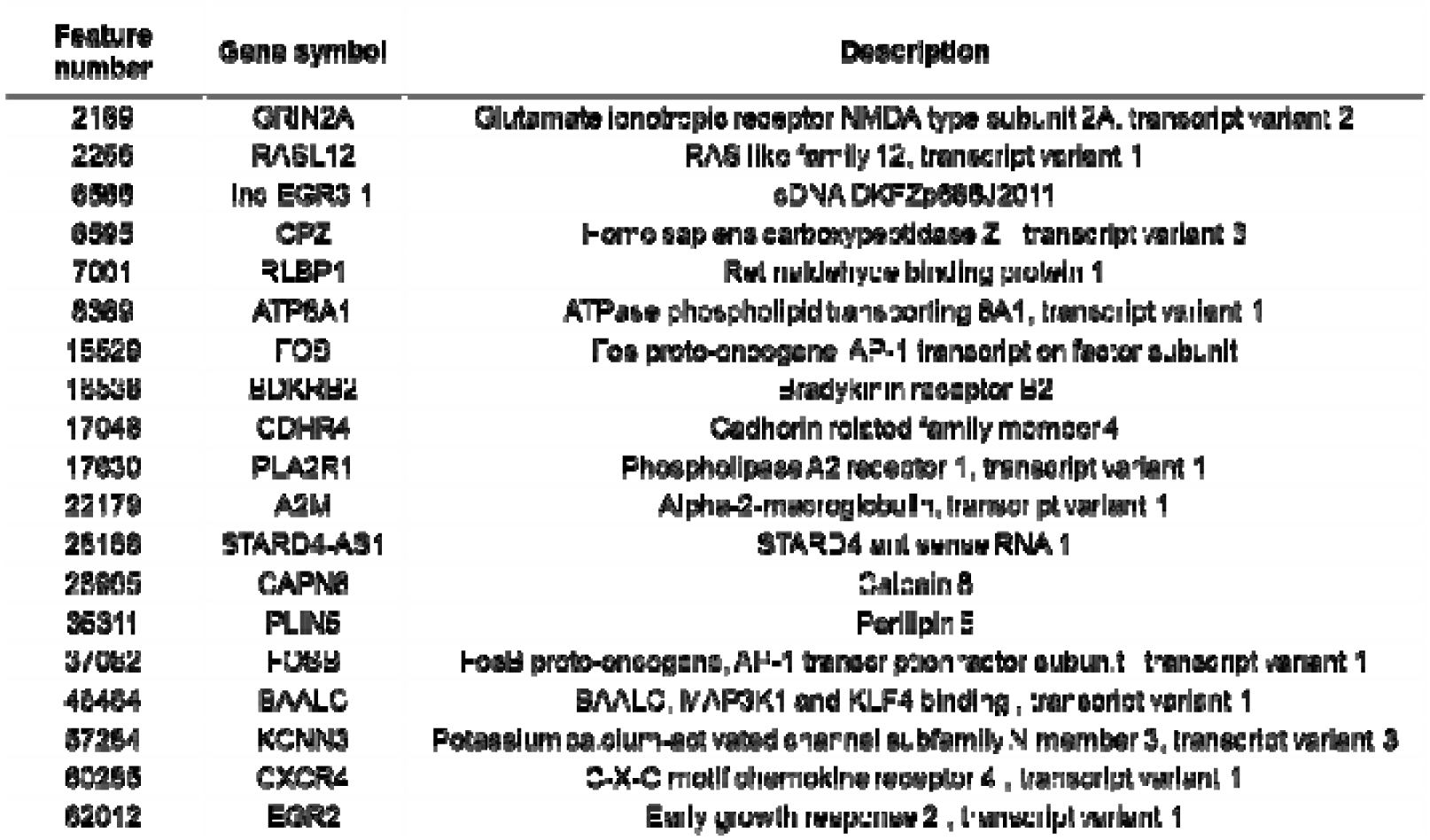

### Analysis of brain invasion cases

In SF4068 meningioma cells with (SF4068 SCN) or without (SF4068 SCC) NF2 mutations^19^, the SF4068 SCN cells exhibited higher expression of stemness markers and CXCR4 compared to the SF4068 SCC cells. Furthermore, this expression was further enhanced in SF4068 SCN cells cultured on hydrogels (Fig. *5A, B*), indicating that CXCR4 levels are elevated in cases harboring NF2 deletions. CXCR4 immunohistochemistry might predict prognosis of meningioma (Fig *5C*).

**Fig. 5.**
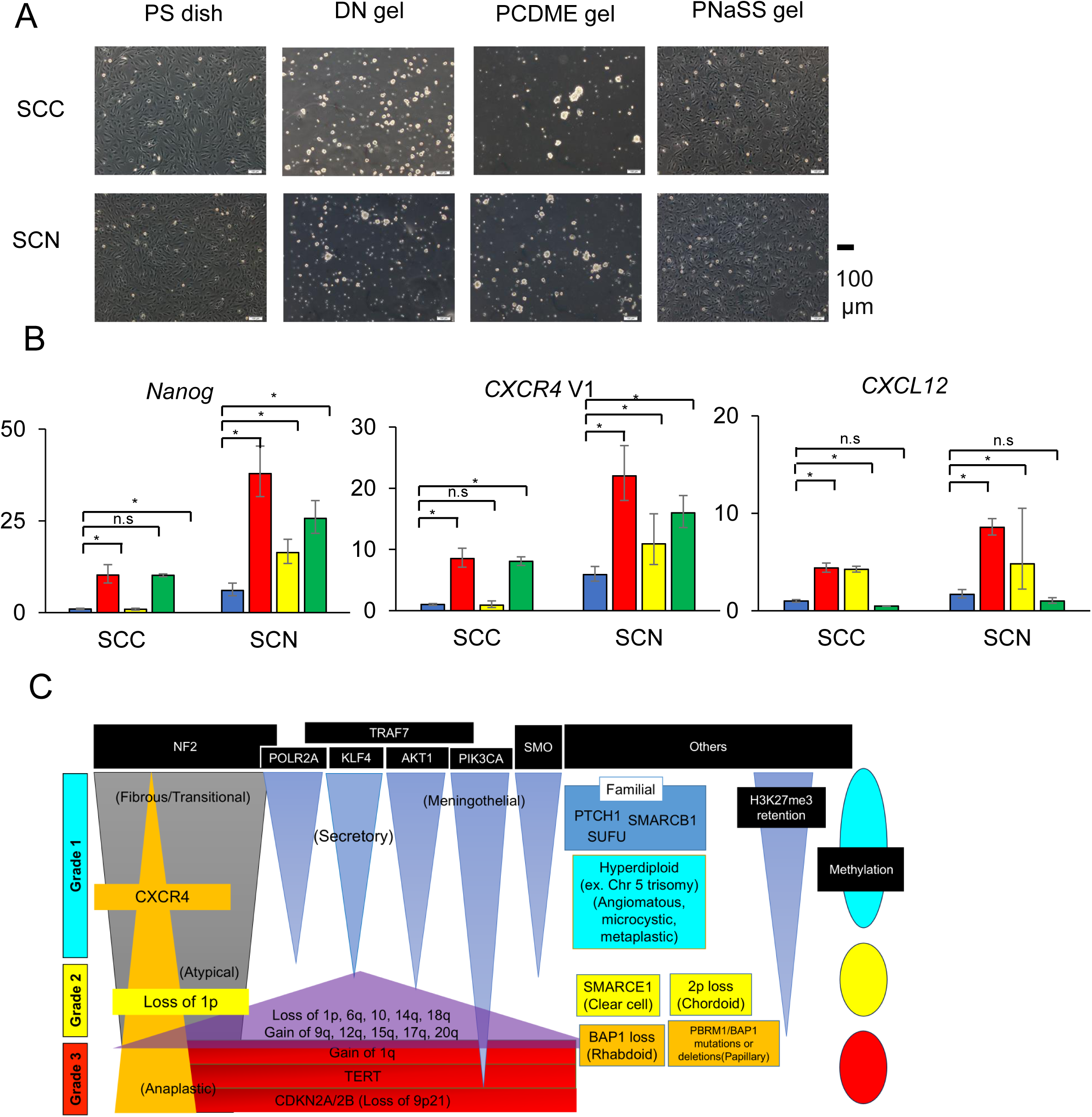
Role of CXCR4 in NF2 deletion meningioma. A. DN gel and PCDME gel induced cell aggregation and forming spherical structures in two cell line (SF4068SCC, SF4068SCN). B. Real-time PCR analysis of the mRNA levels of CXCR4 in two cells (SF4068SCC, SF4068SCN) on DN gel, PCDME gel and PNaSS gel. The data are presented as the mean ± SD of three same experiments in triplicate, and represented using two-sided unequal variance t-tests in comparison with the expression of PS. * P < 0.01 vs. PS. C. Molecular and histological features of meningioma including CXCR4 expression

### Analysis of brain invasion cases

CXCR4 expression is detected in invasion front of meningioma tissues (Fig6. *A*). In case which is scored 3+, CXCR4 positive area extends from the center of the tumor to the invasion front (Fig6. *B*). In cases with both brain invasion and brain adhesion, the part of invasion front was positive for CXCR4, whereas the part of brain adhesion indicated negative for CXCR4 (Fig6. *C*). In brain invasion cases, CXCR4 is positive in 12 of 14 cases and

**Fig. 6.**
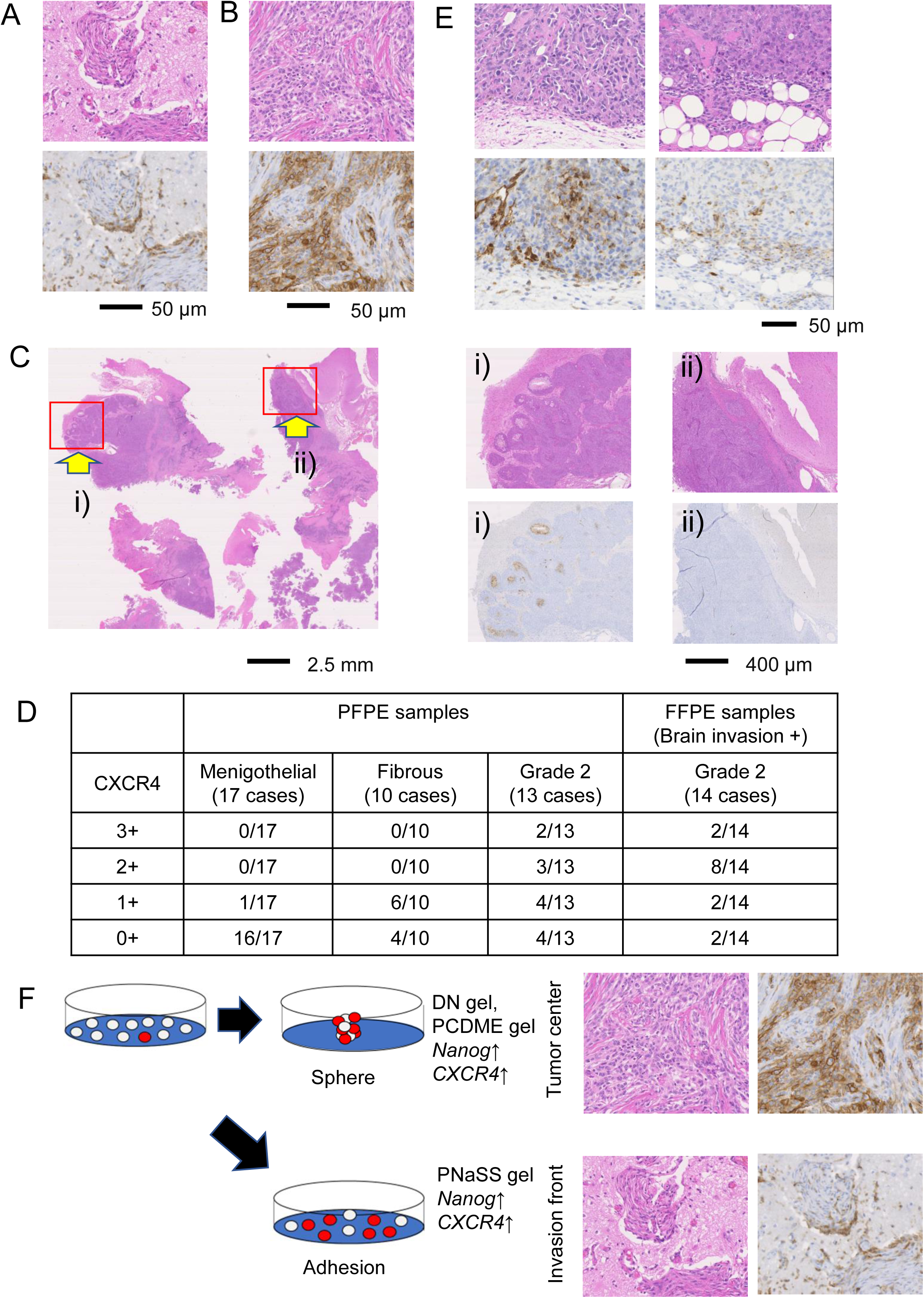
Pathological specimen: Brain invasion. A, CXCR4 expression was observed in invasion front in human meningioma specimen in 13/14 cases. B, In 3+ case, CXCR4 was expressed in the tumor center and invasive front**. C,** Comparison of CXCR4 expression in brain invasion area and brain adhesion area. B, Comparison of WHO grade 1 and WHO grade 2 of CXCR4 positive cell. D, Representative picture of venous invasion. E, CXCR4 immunohistochemistry: CXCR4 was expressed in the tumor center and invasive front. F, Schematic diagram of reprogramming *via* hydrogel in meningiomas

In some cases, it is difficult to determine the presence or absence of brain invasion and adhesion to the brain. We demonstrate CXCR4 might be assistant to diagnose brain invasion (Fig6. *E*).

## Discussion

We recently reported novel method of rapidly inducing CSCs via Hydrogel Activated-reprogramming phenomenon ^12–15^. Here, this study revealed that the PCDME gel and the PNaSS gel in addition to the DN gel were able to induce stemness at the genetic level in meningioma cell lines and primary meningioma cells.

In microarray analysis, 20 candidate stemness genes were selected, which were upregulated 4 times or more by under all conditions using hydrogels compared to the PS dish as control (Fig. 2*D*). CXCR4 has been reported to be associated with tumor proliferative potential in glioblastoma ^21^, and with proliferative and metastatic potential in head and neck cancers ^22^. Furthermore, it has been reported to contribute to poor prognosis in ovarian cancer and osteosarcoma by acting on the tumor microenvironment ^23^ ^24^. Our previous studies have reported that CXCR4 was associated with a poor prognosis in synovial sarcoma^25^. In meningioma, it is known that CXCL12, a ligand for CXCR4, is involved in the acquisition of a stem cell signature^26^. Other than CXCR4, it has also been reported that FOS enhanced the tumorigenicity in osteosarcoma cells ^27^. Grin2a was upregulated in neural stem cells ^28^, and increased in sphere-formed cells of poorly differentiated ovarian cancer ^29^. BAALC has demonstrated to be related to worse prognosis in acute myeloid leukemia with co-expression with MN1^30^. The molecules that were upregulated in the cells on the hydrogel have been associated with poor prognosis in clinical samples.

CXCR4 is a 7-transmembrane G protein-conjugated receptor and is known to be expressed in bone marrow progenitor cells, epithelial cells, microglia, blood cell progenitor cells, lymphocytes, fibroblasts, etc. in normal tissues ^31^. As mentioned above, CXCR4 has been reported to be involved in tumor survival, proliferation, angiogenesis, and metastasis, and it has also been reported to be involved in the interaction between tumor and its surrounding tumor microenvironment *via* the CXCL12-CXCR4 axis. ^32^ ^33^. Indeed, when the recombinant protein of SDF-1 (CXCL12) was added to the cells on the PS dish, an increase in the expression levels of *CXCR4* and the CSC markers were observed (Fig 2*C*).

In xenograft model, it was proved that the tumorigenicity was enhanced in the meningioma cells cultured on the PNaSS gel (Fig. 3*A*), which is one piece of data showing that CSCs have been included. In addition, the formation of satellite nodules and increased venous invasion (Fig. 3*B*, 3*C*) shows that cells cultured on the PNaSS gel might acquire higher infiltration capacity than those than in normal culture. Although the cells on the PNaSS gel did not exhibit a sphere-like morphology, the expression levels of the stem cell markers were increased. Therefore, it was considered that not only the cellular morphological change but also the specific stimulation from the hydrogels induced the cancer stem cell property. In immunohistochemistry, CXCR4 was expressed not only in tumor center but also in invasion front, which indicates that CXCR4 was expressed in an infiltrating cell population. These results suggest that meningioma stem cells exist in the hypoxia niche simulated by the cells cultured on PCDME gel and DN gel at the center of the tumor and in the invasive niche simulated by the cells cultured on PNaSS gel at the tumor infiltration site, and that the distribution of meningioma stem cells is clarified by CXCR4 immunohistochemistry (Fig. 6 *E*).

In some cases, it can be challenging to distinguish between invasion and adhesion of the tumor to brain tissue during meningioma diagnosis. This difficulty arises because, even when the tumor is adhered to the brain parenchyma on histological sections, determining whether this represents true infiltration or merely adhesion often varies among pathologists. However, distinguishing between CNS WHO Grade 1 and Grade 2 meningiomas depends critically on identifying brain infiltration. In this study, a positive CXCR4 staining was observed at the invasive front in cases with confirmed brain infiltration, while no CXCR4 expression was detected in areas where the tumor adhered to the brain parenchyma. These findings suggest that CXCR4 immunostaining may be a useful tool for detecting brain invasion.

While molecular analysis is now required to diagnose meningiomas from WHO 5^th^ edition^2^, genetic analysis is expensive and can only be performed at a limited number of facilities. c-impact Now Update 8 recommend the definition of the cases in which molecular analysis should be used in conjunction with morphological diagnosis ^34^. Molecular analysis is required for cases that require differentiation between CNS WHO grade 1 and CNS WHO grade 2 and 1p and 22q deletion assigned to CNS WHO grade 2. Meningiomas with 1p deletion without 22q deletion were very rare^35^, so predicting 22q deletion can avoid unnecessary molecular analysis.

It is difficult to detect mutations of NF2 without comprehensive analysis because the NF2 gene is huge and there are no hot spots for mutations ^36^ ^37^. In our study, CXCR4 immunohistochemistry was negative in cases with 22q deletion, which is expected to help avoid unnecessary molecular analysis in obvious CNS WHO grade 1 meningiomas. It is not realistic to apply molecular diagnosis to all brain-invasive otherwise benign (BIOB) BIOB meningiomas due to the cost, but immunohistochemistry for CXCR4, which can be a surrogate marker for definitive brain invasion and NF2 deletion or mutation, may be useful for diagnosis of meningioma (Fig. *5E*).

AMD3100, T22, T140, Fc131 are known as inhibitors of CXCR4^38^ ^39^ ^40^ ^41^. Our previous studies have reported that AMD3100 has an *in vitro* tumor growth inhibitory effect on synovial sarcoma ^25^. In this study, AMD3100 was added to the cells on the PS dish and the DN gel, but neither morphological changes nor suppression of stemness marker was observed. We next used Fc131 treatment, but the expression levels of CSC markers were partially suppressed. (Supplemental Fig *1A, 1B*). Stimulation with the recombinant protein of CXCL12 increased the expression level of CXCR4 (Fig *2A*), suggesting that CXCR4-CXCL12 axis may be involved in the acquisition of cancer stem cell properties.

We induced meningioma stem cell signature *via* HARP phenomenon and identified CXCR4 that was biologically and diagnostically significant.

## Material and Methods

### Synthesis of hydrogels

We focused on poly-2-acrylamido-2-methylpropanesulfonic acid (PAMPS) gel, PAMPS/PDMAAm DN gel, Poly N-(carboxymethyl)-N,N′-dimethyl-2-(methacryloyloxy) ethanaminium, inner salt (PCDME) gel, and poly sodium p-styrene sulfonate (PNaSS) gel^16,17,42^. The chemical formula for each gel is shown in (Fig. 1A).

### Cell lines

A human malignant meningioma cell line, HKBMM, obtained from the Riken Cell Bank (Tsukuba, Japan), was cultivated in DMEM/Nutrient Mixture F-12 Ham (Sigma-Aldrich, St. Louis, MO), 15% fetal bovine serum (FBS), penicillin (100 mg/mL), and streptomycin (100 mg/mL). Primary culture cells of meningioma established from surgical specimens ^43^ (WY02, WY08, and WY09) and SF4068SCC and SF4068 SCN obtained from Department of Neuropathology, Otto von Guericke University Magdeburg^19^, were cultured in Gibco™ Ham’s F-12K (Kaighn’s) Medium (reference number 21127022) with 15% fetal bovine serum (FBS). The cells were cultured on a Polystyrene (PS) dish, and a cut-out DN gel, PCDME gel, and PNaSS gel on the bottom of a PS dish.

### RNA extraction and Real-time PCR analysis

Total RNAs of the meningioma cells were extracted using the RNeasy Mini kit (Qiagen, Valencia, CA). cDNA was synthesized using Superscript VILO (Invitrogen, Carlsbad, CA), and Real-time PCR was conducted using SYBR Green PCR Master Mix (Qiagen) and performed on the StepOne real-time PCR system (Applied Biosystems, Foster City, CA). The sequences of the primers are listed in Supplementary Table 1. The relative expression levels of total RNA were normalized to *glyceraldehyde 3-phosphate dehydrogenase* (*GAPDH*).

### Microarray analysis

The WY08 and SY09 cells were cultured for 3 days on PS dish, DN gels, PCDME gel, and for 7 days on PNaSS gel, and total RNA was extracted. Microarray analysis was performed using SurePrint G3 Human Gene Expression 8 × 60K v3.0 (Agilent Array). The data were standardized and analyzed using the process established by the MIAME (Minimum Information About a Microarray Experiment) guidelines. Gene expression levels were analyzed by comparing expression on DN gel, PCDME gel and PNaSS gel with those on PS dish. Hierarchical clustering analysis and heatmap visualization was performed using JMP Pro 14.0.0 (SAS Institute Inc., Cary, NC, USA).

### Patients and specimens

The meningioma resection specimens were obtained from 40 patients who were diagnosed between 1998 and 2012 in Department of Cancer Pathology, Hokkaido University, Sapporo, Japan. This study was approved by the Institutional Review Board of the Hokkaido University, School of Advanced Medicine. Informed consent was obtained from each patient in accordance with the Ethics Committees Guidelines for our institution (Number 17-021).

### Immunohistochemistry

Immunohistochemistry (IHC) for CXCR4 was performed using meningioma 14 formalin-fixed paraffin-embedded (FFPE) meningioma tissues and 36 PAXgene Tissue fixed, paraffin-embedded (PFPE) meningioma tissues. Segment of the comment Hokkaido university. The tissue sections were incubated with 1% bovine serum albumin (BSA) for 10 min at room temperature for preventing non-specific binding of antibodies. After washing, the sections were incubated with anti-CXCR4 antibody (ab124824, abcam) at 100× dilution overnight at 4°C. The subsequent processes were carried out according to the manufacturer’s instructions using the Envision FLEX visualization system (Dako; Agilent Technologies, Inc., Santa Clara, CA).

### Xenograft model

Six-to seven-weeks old female SCID mice, Balb/cA Jcl nu/nu (Clea Japan, Inc., Tokyo, Japan) were used for a xenograft model. WY08 cells seeded on PS dish or on PNaSS gel at a concentration of 1.0 × 10^5^/ml, after 1 week, the 1.0 × 10^3^, 1.0 × 10^4^, and 1.0×10^5^ cells (each n=4) were injected subcutaneously into both abdomen of mice. After 11 weeks, mice were sacrificed and tumors were removed from subcutaneous tissue, weighed and sized, and FFPE tissues were constructed, and histological evaluation was performed. Experiments were conducted in compliance with the guidelines of Hokkaido University Manual for Implementing Animal Experimentation with the approval of Institutional Animal Care and Use Committee at Hokkaido University Graduate School of Medicine (Number 17-0061).

### Statistical analyses

We used JMP Pro 14.0.0 (SAS Institute Inc., Cary, NC, USA) for statistical processing. Data are presented as mean ± SD. Comparisons of continuous data between two groups were performed using Student’s unpaired t-test.

## Data Availability

All study data are included in the article or *SI Appendix*.

## ACKNOWLEDGMENTS

We thank Ms. Yukiko Hane (Hokkaido University) for gel synthesis, Ms. Mieko Hishikawa (Hokkaido University) for immunohistochemistry. This work was supported in part by Grants-in-Aid from the Ministry of Education, Culture, Sports, Science, and Technology (19H01171 for S.T.), from the Japanese Society for the Promotion of Science, and from the Ministry of Health, Labor, and Welfare of Japan, as well as by a grant from the Japanese Science and Technology Agency. Institute for Chemical Reaction Design and Discovery (ICReDD) was established by World Premier International Research Initiative (WPI), MEXT, Japan. In addition, this research was supported by Global Station for Soft Matter, a project of Global Institution for Collaborative Research and Education at Hokkaido University.

---

**Supplemental figure 1.**
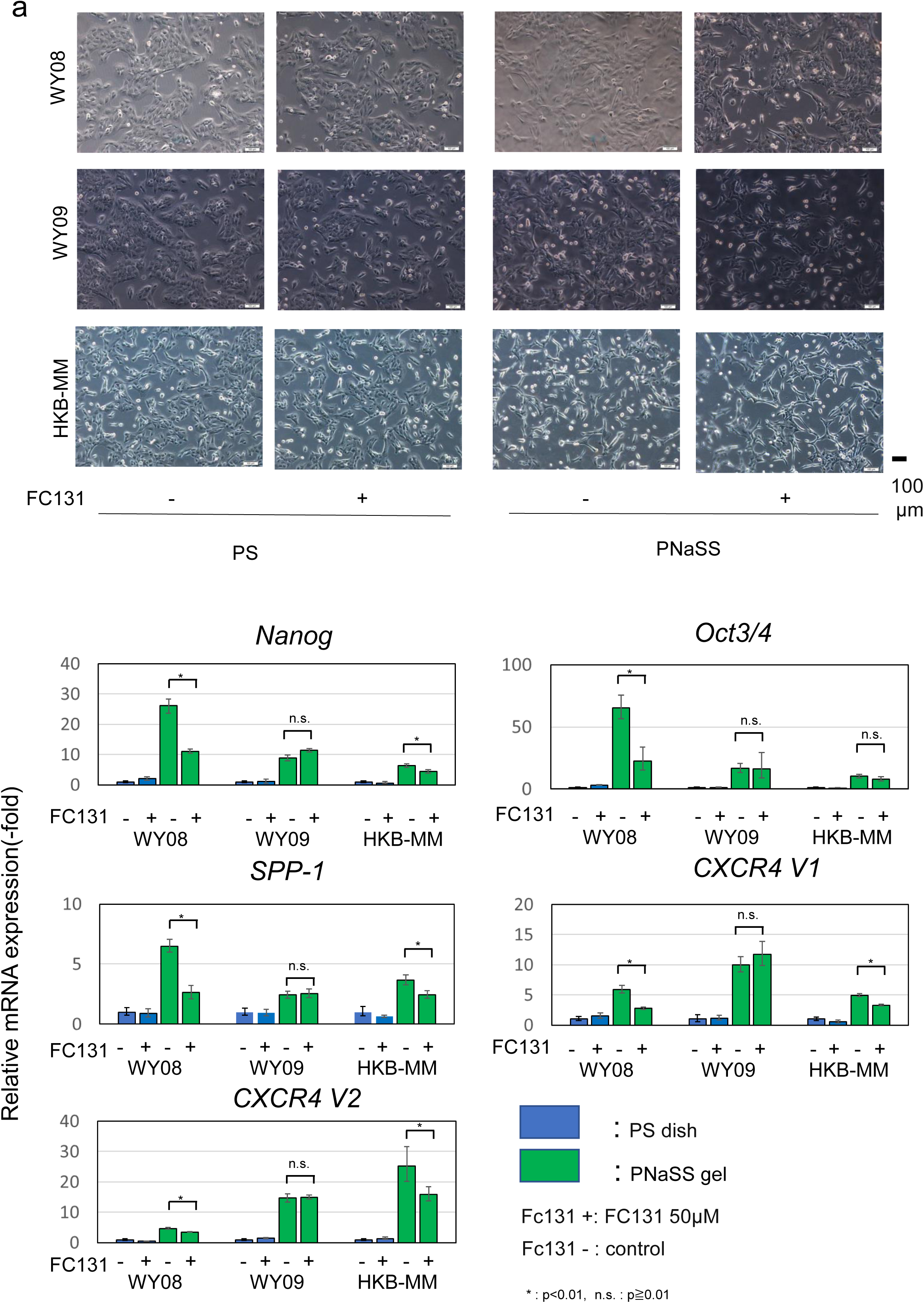
Treatment with Fc131 suppressed stemness induction of meningioma on PNaSS gel. **a**, Representative photographs of WY08, WY09, HKB-MM cells cultured on PS dish and PNaSS gel with or without 50 µM Fc131(CXCR4 inhibitor). **b,** The expression level of *Nanog, Oct3/4, SPP-1, CXCR4(V1, V2)* was depressed by FC131 on PNaSS gel.

